# Multimodal cellular context enables molecular phenotyping from single-molecule trajectories

**DOI:** 10.64898/2026.11.05.764606

**Authors:** Taras Redchuk, Lea Rehm, Yuan Sui, Jarno Mäkelä

## Abstract

Single-molecule tracking (SMT) provides direct measurements of molecular dynamics in living cells, but motion alone rarely specifies the underlying biological process, without accounting for the cellular environment in which that motion occurs. Here, we present MIST (Multimodal Interpretation of Single-molecule Trajectories), a framework that brings the paradigm of image-based profiling to single molecules by integrating trajectory dynamics, emitter properties and cellular context into a common, reusable phenotype space. Using a reference panel of nine proteins in live *Escherichia coli*, we show that multimodal context more than doubled protein-specific information relative to trajectory features alone, a gap that learned trajectory representations did not close. MIST allowed de novo identification of shared transcription-translation phenotype at the nucleoid periphery. A fixed reference space detected effects of previously uncharacterized MukB ATPase mutations without retraining. MIST provides an interpretable and extensible framework for comparing proteins, mutants and conditions within a common space of molecular phenotypes.

## Introduction

Single-molecule tracking (SMT) measures how individual molecules move in their local environment and is widely used to study molecular behaviour in living cells^1–4^. A broad range of approaches has been developed to describe molecular motion^5^, quantify binding^6,7^, resolve discrete diffusive states^8–11^ and probe the heterogeneity of protein behaviour^12–14^. Diffusional fingerprinting combines engineered trajectory features with hidden Markov model-derived states^15^, and machine learning extracts additional information from trajectories, with models assigning states to individual trajectory steps^16,17^.

Despite this, the biological interpretation of SMT data often remains ambiguous, because motion alone rarely specifies the process that produces it. Different modes of confinement and cellular environments can produce similar behaviour^18,19^, whereas a protein may switch between states, forming mixtures of populations that are difficult to separate^20,21^. Molecular motion also depends on the physiological state of the cell, as cell-cycle progression, stress and nutrient availability alter the intracellular environment, including cytoplasmic viscosity and macromolecular crowding^22,23^. Because such factors are rarely quantified independently from the trajectories, their effects on molecular motion are usually not accounted for. More sophisticated models cannot resolve such ambiguity, because the distinguishing information is not often contained in the trajectories.

Image-based profiling addresses related challenges at the cellular level. Cell Painting combines stains of several cellular compartments into unbiased, high-dimensional morphological profiles^24,25^, and general organelle morphology carries information comparable to that of omics measurements^26^. Proteome-wide localization atlases apply a single representation across protein libraries to reveal functional relationships^27–29^, and cytological profiling classifies antibiotic mechanisms of action in bacteria ^30^ and maps gene function from cell morphology and cell-cycle phenotypes^31,32^. Each acquisition modality captures only part of the biological process under study, and integrating several modalities into comprehensive, reusable representations remains a challenge for both biology and computer science^33^, addressed in single-cell genomics by dedicated integration frameworks^34,35^.

SMT and image-based profiling therefore have complementary blind spots. Profiling yields reusable phenotypes without molecular resolution, whereas SMT measures molecular behaviour directly but, to our knowledge, has not been combined with the profiling concept. Recent SMT methods have begun to incorporate photometric properties and subcellular localization^17^, but these focuses to analyse one protein at a time, rather than to construct a comprehensive representation. What is missing is a modular framework that describes each trajectory in its cellular context, integrates SMT with complementary imaging modalities and can be reused across proteins, perturbations and biological questions. Bacteria provide a suitable test system for such framework: their cells are compact yet spatially organized, with the membrane, cell poles and nucleoid defining distinct subcellular environments that can be resolved by light microscopy^36,37^.

Here, we introduce MIST (Multimodal Interpretation of Single-molecule Trajectories), a framework that extends SMT from the analysis of molecular motion to context-aware molecular phenotyping in living cells. MIST combines trajectory dynamics with emitter properties and independent measurements of cell and nucleoid organization in a common phenotype space, while keeping physically interpretable features at its core. Using a reference panel of nine proteins in live *Escherichia coli*, we quantify the information contributed by each feature family and show that multimodal context more than doubles protein-specific information relative to trajectory features alone, exceeding what learned trajectory representations achieve. Deep-learning-derived features are nevertheless incorporated as an optional module. MIST resolves proteins within families of freely diffusing, membrane-associated and nucleoid-associated phenotypes, and identifies de novo a phenotype jointly enriched in RNA polymerase and ribosome trajectories, consistent with a spatial interface between transcription and translation. Finally, a fixed MIST reference space detects redistribution of molecular phenotypes in previously unseen MukB mutants, including previously uncharacterized substitutions. Together, these results establish multimodal molecular phenotyping as a general approach that brings the profiling paradigm to single molecules.

## Results

### A multimodal single-molecule reference dataset

Proteins perform a wide range of roles in the cell and thus occupy a broad and heterogeneous behavioural space. To map this variability in living cells, we assembled a reference panel of proteins tagged with HaloTag expressed from their native chromosomal locus covering distinct functions associated with particular subcellular localizations or modes of action (Fig. 1A; Table 1). The panel comprises freely diffusing cytoplasmic proteins (HaloTag alone; metabolic enzyme, MetE), membrane-associated proteins (polar, inner membrane, TsR; non-polar, outer membrane, LptD), nucleoid-associated proteins (histone-like, H-NS; structural maintenance of chromosomes (SMC) complex subunits, MukB and MukE) and components of large macromolecular assemblies (RNA polymerase, RpoC; ribosomal 30S subunit, RpsB). MukB-HaloTag was also expressed in a second genetic background (AB1157) containing multiple auxotrophies and sugar utilization defects, which perturbs growth, cell size, and replication state among others, providing a context perturbation of an otherwise identical protein. HaloTag was fluorescently labelled using Janelia Fluor 554 dye (JFX554) and single-molecule trajectories were collected in live cells for each protein of interest. All together, we analysed 7.8M protein localizations forming 300k trajectories in 5700 cells across 3 biological replicates. In addition to single-molecule trajectories, we imaged each cell with bright field and with the DNA stain SYTO9, which allowed us to segment individual cells, derive their geometry and a cell-centred coordinate system, and map nucleoid organization, respectively, so that every trajectory could be positioned relative to the cell boundary and the nucleoid (Fig. 1B). Together, these measurements form a multimodal reference dataset (MISTD v0.1), which is publicly available, in which each single-molecule trajectory is linked to orthogonal descriptors of its cellular context (Fig. 1B; Data availability).

**Fig. 1.**
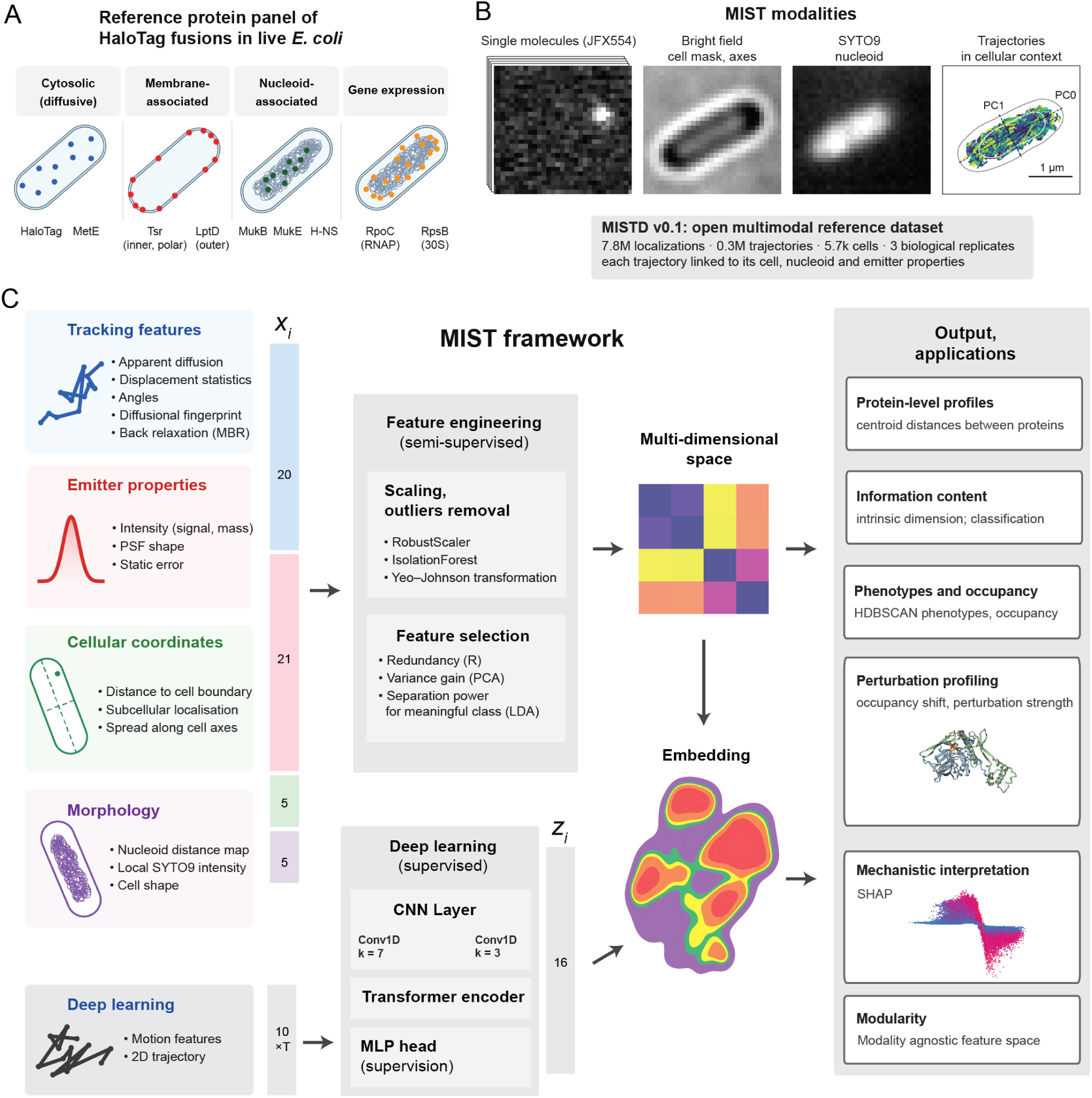
MIST framework. **A**, Reference panel of nine proteins imaged in live Escherichia coli, grouped by expected behaviour: cytosolic, freely diffusing (HaloTag alone, MetE); membrane-associated (Tsr, inner membrane, polar; LptD, outer membrane); nucleoid-associated (MukB, MukE, H-NS); and gene expression (RpoC, β′ subunit of RNA polymerase; RpsB, 30S ribosomal protein S2). **B**, Imaging modalities used in MIST. Single molecules panel (left) shows 1 frame from the timelapse. Tracking panel shows all detected tracks from the selected cell. Scale bar, 1 µm. **C**, Analysis pipeline. Input data is shown on the left, *x*_i_ denotes the feature vector used in the MIST representation. *z*_i_ denotes latent features as returned by DL module (CNN Transformer).

**Table 1.** Proteins used in this study.

| Strain name | Gene | Protein name | Function | Back-ground |
|---|---|---|---|---|
| CJW7519 | *rpoC* | RNA polymerase | DNA-binding | MG1655 |
| JML_Ec_136 | *metE* | MetE | non-interacting | MG1655 |
| CJW7528 | *rpsB* | Small ribosomal subunit (30S) | Ribosome | MG1655 |
| JML_Ec_173 | *tsr* | Methyl-accepting chemotaxis protein 1 | Membrane, at poles | MG1655 |
| LT18 | *lptD* | Lipopolysaccharide (LPS)-assembly protein LptD | Outer membrane | MG1655 |
| JML_Ec_15 | *hns* | DNA-binding protein H-NS | Chromosome organization | MG1655 |
| JML_Ec_01 | *mukB* | Chromosome partition protein MukB | DNA-binding | MG1655 |
| JML_Ec_59 | *halotag* | cytosolic HaloTag | non-interacting | MG1655 |
| JM48 | *mukE* | Chromosome partition protein MukE | DNA-binding | AB1157 |
| JM41 | *mukB* | Chromosome partition protein MukB | DNA-binding | AB1157 |
| JM65 | *mukB(E1407Q)* | Chromosome partition protein MukB mutant | DNA-binding (altered) | AB1157 |
| JM66 | *mukB(D1406A)* | Chromosome partition protein MukB mutant | DNA-binding (altered) | AB1157 |
| JM67 | mukB(Walker B/D-loop(D1412E)) | Chromosome partition protein MukB mutant | DNA-binding (altered) | AB1157 |
| JM68 | mukB(Walker B/D-loop(D1412A)) | Chromosome partition protein MukB mutant | DNA-binding (altered) | AB1157 |
| JM71 | mukB (Signature(L1365V, G1368S)) | Chromosome partition protein MukB mutant | DNA-binding (altered) | AB1157 |

### The MIST framework

Single-molecule trajectories provide a direct description of molecular motion but do not uniquely specify the biological process underlying that motion. We therefore sought to embed individual trajectories from all proteins and experiments in a common representation, in which molecular behaviours could be compared directly at single-trajectory resolution. For example, cellular context distinguishes slow motion at the cell periphery from slow motion within the nucleoid, separating membrane-associated and chromosome-associated behaviours that can overlap strongly in diffusion space. To this end, we represented each trajectory at four levels of information (Fig. 1c): (1) track-level features (20 features), including the apparent diffusion coefficient (D_app_) derived features, track length, displacement angles, diffusional fingerprinting features (^15^, excluding hidden Markov-derived features), and mean back relaxation (MBR)^38^; (2) emitter properties (21 features) obtained during localization, including signal intensity and spot morphology, which describe the detected particle independently of its reconstructed coordinates; (3) cellular geometry (5 features), obtained by registering each trajectory to its bright-field-derived cell mask, which defines a normalized cellular coordinate system and spatial descriptors such as distance to the cell boundary and position along the cell axes; and (4) cell morphology and nucleoid organization (5 features), which describe each trajectory relative not only to the cell boundary but also to the chromosome-containing region of the cell.

Because the different modalities yield features with different scales, redundancy and sensitivity to experimental variation, we processed them in three steps before integration. First, features were transformed and standardized before integration. Second, we removed highly correlated and weakly informative features using principal component analysis (PCA). Third, we used linear discrimination analysis (LDA) to prioritize features that separated proteins more strongly than biological replicates. This procedure retained 31 of the 51 input features, which we use for all subsequent analyses.

The resulting feature vectors define a high-dimensional molecular phenotype space in which each trajectory is a single point, determined jointly by trajectory dynamics, emitter properties, and cellular context, and in which proximity reflects similarity across multiple dimensions of molecular behaviour. Each protein is therefore represented by a distribution of points in this space. In image-based proteomics, distances between protein-level profiles have been used to infer functional relationships and even detect protein-protein interaction^29,39^. To obtain a coarse summary of each protein, we calculated the centroid of each protein’s feature vectors and quantified pairwise Euclidean distances between centroids (Fig. 2A).

**Fig. 2.**
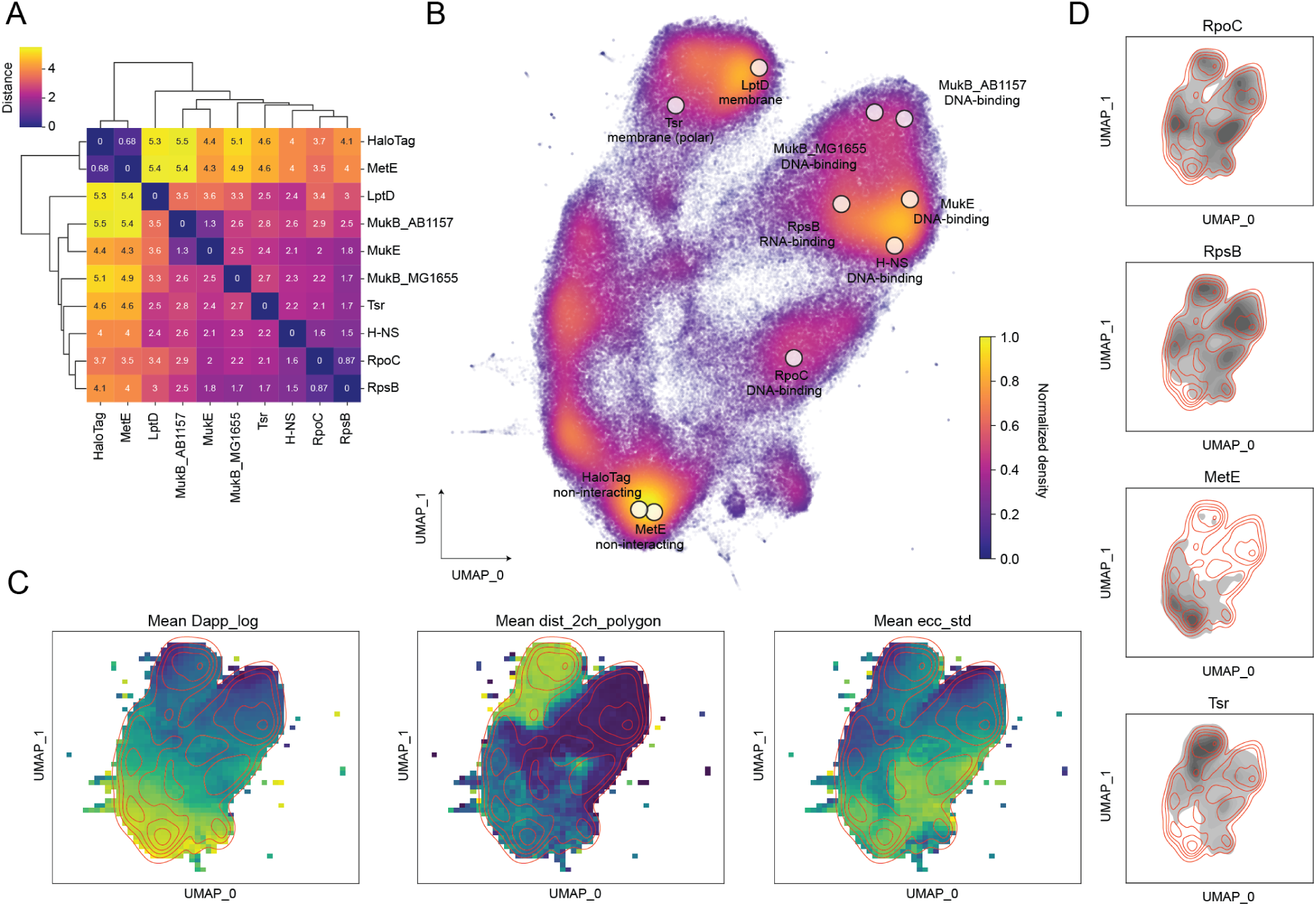
MIST organizes single-molecule trajectories into a shared molecular phenotype space. **A**, Euclidean distances between protein centroids (mean feature vectors) in the multidimensional MIST feature space, with average-linkage hierarchical clustering. Functionally related proteins are close: HaloTag and MetE form the closest pair (0.68), and the MukBEF components MukB and MukE are 1.3 units apart. **B**, UMAP projection of trajectories coloured by normalized point density. Circles mark the position of each protein’s maximal density and are labelled with its expected function. **C**, Mean D_app_ (apparent diffusion coefficient, left), minimal distance to the nucleoid outline; 0 inside the nucleoid polygon (middle) and standard deviation of the emitter eccentricity (right), binned and mapped to the UMAP embedding in B. Red lines, isolines of the reference trajectory density (identical in C and D). Selected feature maps are shown, all feature maps are shown in Supplementary Fig. 1. **D**, Trajectory density (grey) of RpoC, RpsB, MetE and Tsr over the reference isolines. MetE is confined mostly to the high-mobility region and Tsr to the peripheral low-mobility region, whereas RpoC and RpsB populate several regions, consistent with heterogeneous single-molecule behaviour.

Proteins with related molecular behaviours occupied nearby position in this space (Fig. 2A): HaloTag and MetE, both dominated by free diffusion in the cytoplasm, formed the closest protein pair (distance 0.68); MukE, being part of the MukBEF complex, was close to MukB in the same AB1157 strain background (1.3) but further from MukB in MG1655 (2.5), consistent with the differences between the two backgrounds. Thus, protein-level distance reflects both molecular behaviour and the cellular context in which it occurs.

### Mapping the molecular phenotype landscape with MIST

Protein-level centroids necessarily average over the heterogeneity captured by single-molecule measurements. Proteins dominated by a single mode of behaviour are expected to form relatively compact distributions, while proteins that partition between functional states, interaction partners, or intracellular environments, should occupy multiple regions of the phenotype space. We next examined the representation at the level of individual trajectories.

To visualize the high-dimensional phenotype space, we projected the feature vectors into two dimensions using UMAP^40^ (Fig. 2B). Each trajectory corresponds to one point in the embedding, and its local neighbourhood reflects similarity in the underlying multimodal representation. The embedding contained several densely populated regions connected by less densely populated areas. Although UMAP coordinates cannot be inverted back to high-dimensional space, mean value of each feature can be calculated across the embedding to generate feature maps (Fig. 2C, Supplementary Fig. 1). Individual features varied across the embedding in distinct ways: D_app_ formed a continuous gradient along the diagonal of the embedding; in contrast, distance to the nucleoid formed two well-separated extrema within the low-D_app_ region, separating slowly moving trajectories far from the nucleoid from those overlapping it; while variability in spot eccentricity followed a pattern distinct from both. Thus, trajectories with similar mobility can occupy distinct regions of the MIST representation once their cellular context is considered. Because no single input feature accounted for this organization, the representation reflects combinations of molecular and cellular properties.

We next asked how individual proteins were distributed across this landscape (Fig. 2D). Proteins with similar expected molecular behaviours populated related regions of the embedding, whereas proteins participating in multiple molecular processes occupied broader portions of it. MetE was strongly enriched in the high-mobility region, consistent with unrestricted cytoplasmic diffusion, whereas the membrane protein Tsr mainly occupied a low-mobility region distant from the nucleoid, consistent with its localization at the cell periphery. In contrast, RpoC and RpsB each extended across several regions, reflecting partitioning of RNA polymerase and ribosomal subunits between active and non-active states. Notably, their distributions were similar, in line with their short centroid distance (Fig. 2A).

Together, these analyses show that MIST organizes single-molecule trajectories according to combinations of molecular dynamics and cellular context, and that functionally related proteins populate similar regions of the resulting landscape. However, these observations are largely descriptive. We therefore asked how much biologically meaningful information the MIST representation retains, and whether cellular context improves discrimination between proteins beyond trajectory dynamics alone.

### Quantifying molecular information in MIST

We quantified how much information each feature family contributes using two complementary measures: an unsupervised estimate of non-redundant information and a supervised measure of protein-specific information.

First, we estimated intrinsic dimension (ID) of the data with TwoNN estimator^41,42^, while sequentially adding the four MIST feature families: track-level features (tracking), emitter properties (Em), cellular coordinate-based features (PC_coord), and cell morphology (Morph). ID reports the number of independent directions of variation needed to describe the data and thus serves a proxy for non-redundant information. To separate new information from redundancy, we compared the ID of each feature set with that of a control in which each feature was shuffled independently across trajectories, which destroys the shared structure between features while preserving their individual distributions (Fig. 3A). In this comparison, points close to the identity line (x=y) correspond to features that vary independently, as expected for noise, while points below it indicate structured, partly redundant variation. Every feature family increased the ID. Adding Em raised the ID from 11.4 to 14.9 and adding PC_coord raised it further to 16.5, in each case by roughly half of the corresponding increase in shuffled ID, indicating that both families contributed new variation that remained structured with the rest of the data. In contrast, Morph increased the ID only marginally (to 16.9), presumably because cell morphology is already partly captured by the cellular coordinates of the emitters.

**Fig. 3.**
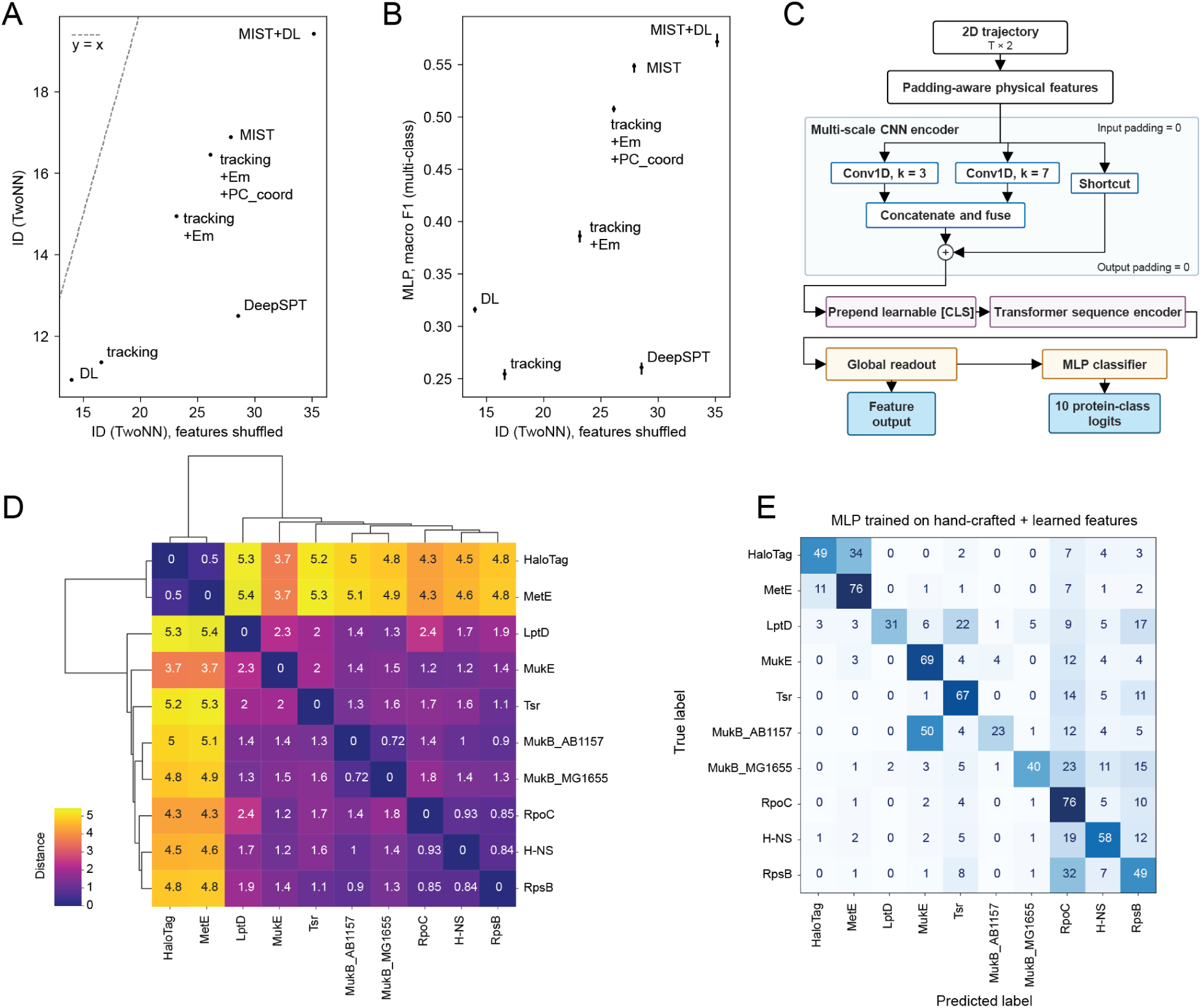
Cellular context adds information that trajectory dynamics alone do not capture. **A**, Intrinsic dimension (ID; TwoNN estimator) of each feature set versus its ID after independent permutation of every feature, which preserves feature distributions but removes shared structure (horizontal bars, s.d. over 3 permutations). Emitter properties (Em), cell-coordinate features (PC_coord) and morphology (Morph) are added cumulatively to the tracking features; MIST+DL further includes the learned features from C. DeepSPT, DeepSPT-derived features. Distance below the dashed y = x line reflects redundancy among features. **B**, Macro-averaged F1 score of a multilayer perceptron (MLP; three hidden layers of 100 units) classifying the 10 reference strains, versus shuffled ID. Points, mean of 5 random cross-validation splits stratified by protein and biological replicate; vertical bars, min-max range. Emitter and cellular context more than double classification performance relative to tracking features or DeepSPT, whereas higher ID alone does not improve protein discrimination. **C**, Architecture of the physics-informed CNN-Transformer. The global readout gives the 16-dimensional latent features (DL); the network is trained through an MLP head with 10 protein-class outputs. **D**, Euclidean distances between protein centroids in the DL latent space, with hierarchical clustering. **E**, Confusion matrix of an MLP trained on engineered and learned features. Values, percentage of trajectories of each true class assigned to each predicted class; classes are ordered as in d. Misclassification is higher between proteins with similar expected behaviour, such as HaloTag and MetE, or MukB(AB1157) and MukE.

We next evaluated protein-specific information by training a multilayer perceptron (MLP) classifier to predict protein identity from each feature set. Although protein identification is not the intended application of MIST, the reference panel was designed to cover a broad range of molecular behaviours and classification performance therefore reports how much protein-specific information a representation contains. Trajectory features alone yielded a macro-averaged F1 score of 0.25 (chance level, 0.10), reflecting the substantial overlap in motion between many reference proteins (Fig. 3B). Sequential addition of emitter properties (F1 = 0.39), cell coordinates system (F1 = 0.51) and morphology context (F1 = 0.55) progressively improved classification, more than doubling performance relative to trajectory features alone. Comparison of the confusion matrices showed that cellular context mainly improved discrimination between proteins with overlapping mobility but different spatial environments, whereas HaloTag and MetE remained difficult to distinguish. Interestingly, morphology improved classification despite its small ID gain, suggesting that the non-redundant part of morphological information is specifically relevant to protein identity. Taken together, MIST integrates complementary trajectory, emitter, spatial and morphological information into a representation with measurable non-redundant and protein-specific information content.

### Benchmarking against learned trajectory representations

We next asked whether the limited performance of trajectory only features reflected the choice of engineered descriptors rather than the information available in trajectories themselves. To test this, we compared MIST with two learned trajectory representations: the state-of-the-art method DeepSPT^16^, which extracts temporal and diffusional features from particle trajectories, and a supervised deep-learning (DL) model that we trained directly on single-molecule tracks. For the DL model, we developed a physics-informed CNN–Transformer architecture (Fig. 3C), in which one-dimensional convolutional branches capture local motion patterns at different temporal scales and a Transformer encoder integrates information across the full trajectory. The resulting representation was compressed into a 16-dimensional latent space, which we evaluated with the same ID analysis and the same MLP classifier as the engineered features.

Both learned representations performed similarly to engineered tracking features. DeepSPT had a higher ID than either engineered tracking or DL features (12.5 vs. 11.4 and 10.9, respectively; Fig. 3A) but achieved a macro F1 score of only 0.26, close to the tracking baseline (Fig. 3B). DL features performed better (F1 = 0.32), as expected for a representation trained with protein labels, despite having the lowest ID. Thus, greater intrinsic complexity does not necessarily translate into protein discrimination, and all three trajectory representations remained far below the full MIST representation (F1 = 0.55), which more than doubled the performance of DeepSPT.

Distances between protein centroids in the DL latent space (Fig. 3D) showed that the learned representation was biologically structured but compressed relationships that MIST resolved. HaloTag and MetE again formed the closest pair (0.50), whereas RpoC, RpsB and H-NS were all within 0.93 of each other, consistent with similar motion. Notably, the two different strains with MukB were close in DL space (0.72) but well separated in MIST (2.6; Fig. 2A), consistent with their difference being carried by cellular context rather than by MukB dynamics.

Combining DL features with the full MIST representation (MIST+DL) further increased the ID (to 19.4) but improved classification only modestly (F1 = 0.57; Fig. 3B). The remaining misclassifications were largely biologically interpretable (Fig. 3E): HaloTag was frequently assigned as MetE (34%), LptD as Tsr (22%), RpsB as RpoC (32%) and MukB_AB1157 as MukE (50%), pairs that share molecular behaviour, cellular compartment or genetic background. Given the modest gain, the largely unstructured global organization of the DL latent space (Supplementary Fig. 2) and the limited interpretability of learned features, it is unclear how much of the additional information is biologically meaningful. We therefore keep DL as an optional module, which illustrates the modularity of MIST, and use the engineered representation for all subsequent experiments.

### MIST resolves molecular states

The molecular phenotype landscape contained dense regions occupied by different proteins. To quantify the molecular dynamics in these regions, we clustered trajectories in the MIST representation and interpreted the resulting populations using their dynamic and spatial properties and feature attributions.

We used density-based clustering (HDBSCAN)^43^, which identifies recurrent populations without specifying their number or imposing predefined molecular states. After excluding a small cluster enriched for tracking artifacts, the analysis resolved four trajectory populations (P1-P4; Fig. 4A;). Cluster assignments were robust to variation in clustering parameters (Supplementary Fig. 3), indicating that they reflect stable organization of the phenotype space.

**Fig. 4.**
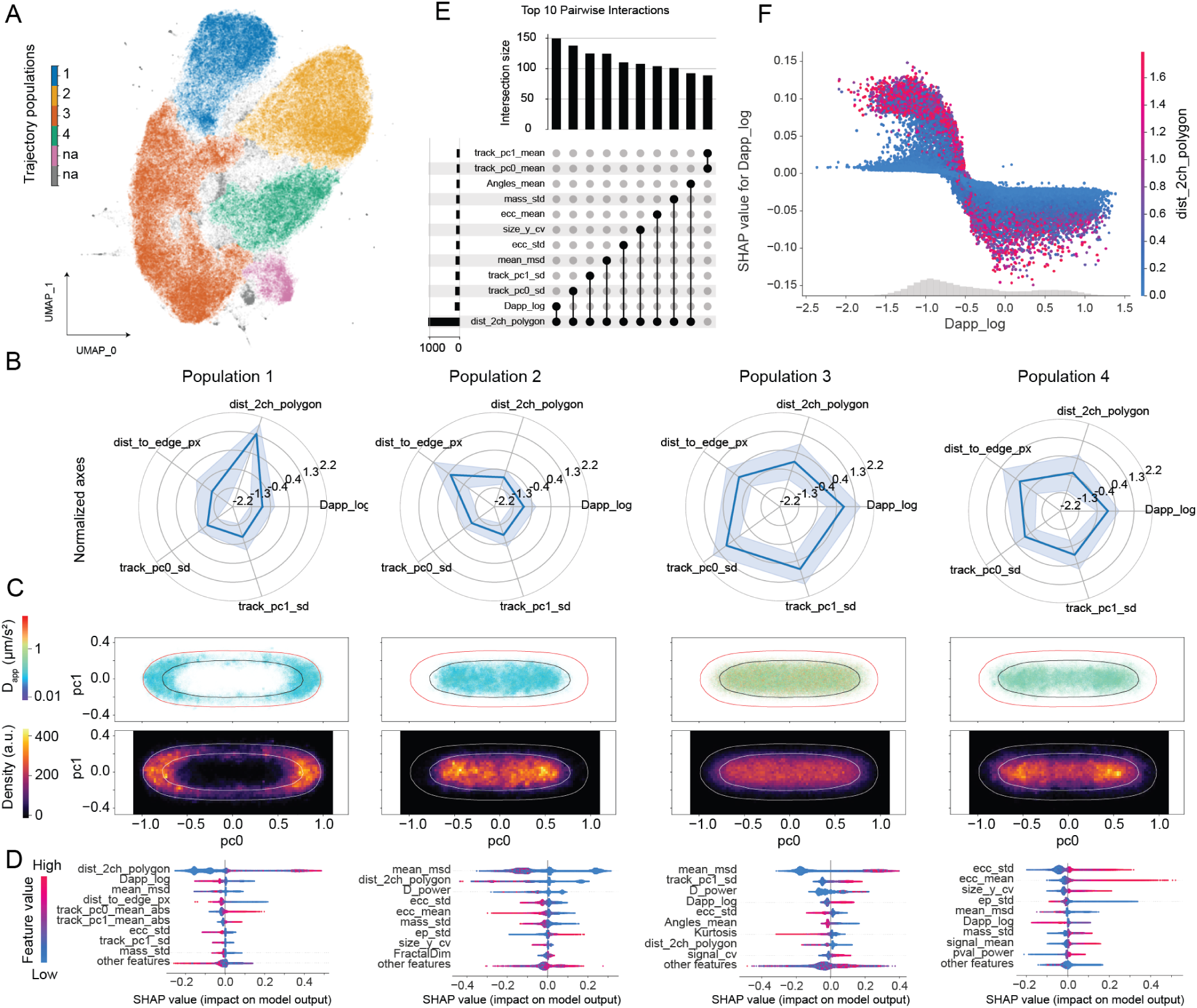
Density-based clustering resolves recurrent molecular phenotypes defined by combinations of motion and cellular context. **A**, Embedding from Fig. 2B coloured by trajectory population as identified by HDBSCAN. na, unassigned trajectories (grey) and cluster enriched in tracking artefacts (pink), both excluded from downstream analyses. **B**, Radar plots of five features for populations 1-4: apparent diffusion coefficient (D_app_), distance to the nucleoid, distance to the cell edge, and trajectory spread along the long and short cell axes. Values are transformed and standardized; line, mean; shaded area, s.d. Descriptive statistics are given in Supplementary Table 1. **C**, Localizations of each population in normalized cell coordinates, coloured by D_app_ (top), and the corresponding localization density (bottom; binned). Lines, dataset-wide outlines of cell masks and SYTO9-stained nucleoids. **D**, SHAP summary plots for one-versus-rest gradient-boosting classifiers (GBC) trained on all MIST features. SHAP values of predicted probability were computed on the 20% held-out trajectories; each dot is one trajectory, coloured by feature value, and the nine highest-ranked features are shown. **E**, The ten strongest pairwise feature interactions for P1, ranked by approximate SHAP interaction strength. Distance to the nucleoid takes part in nine of the ten. **F**, SHAP dependence plot of D_app_ for P1, coloured by distance to the nucleoid; grey histogram, distribution of D_app_. Low mobility favours assignment to P1 mainly for emitters far from the nucleoid, and the effect weakens for emitters near or inside it. Membrane- and nucleoid-associated trajectories thus overlap in mobility and are separated by their cellular context.

Three populations were defined by distinct combinations of mobility and intracellular position (Fig. 4B, C, Supplementary Table 1). P1 population consisted predominantly of slowly moving trajectories (median D_app_ = 0.11 µm²/s) located near the cell periphery and outside the nucleoid (median distance from nucleoid 0.095 µm), consistent with membrane-associated molecules. P2 contained similarly slow trajectories within the nucleoid, with both 25th and 75th percentiles of nucleoid distance inside the nucleoid mask, consistent with nucleoid-associated molecules. P3 was characterized by higher mobility (median D_app_ = 1.6 µm²/s), and its trajectories extended along the long cell axis at least 3-fold more than those of the other populations (SD 0.18 in normalized cell coordinates), consistent with freely diffusing molecules. Notably, P1 and P2 partially overlapped in their diffusional properties and were separated mainly by their position relative to the nucleoid. P4 exhibited a different but less defined combination of features.

To determine systematically which features distinguish the populations, we trained one-versus-rest gradient-boosted classifiers (GBC) on the MIST representation and quantified the contribution and direction of each feature using SHAP (Shapley additive explanations)^44^. Because the value of SHAP lies in capturing nonlinear and conditional feature effects, we first asked whether such effects were present by comparing each GBC with a logistic regression classifier. The GBC outperformed the linear model of all populations, with gains in balanced accuracy varying ranging from 0.8 percentage points (P2) to 8.9 percentage points (P1), indicating that nonlinear feature effects contribute to population identity, most strongly for P1.

SHAP analysis confirmed the mobility and cellular position defined P1-P3 (Fig. 4D) and, in addition, revealed contributions from emitter properties that were not apparent from the population profiles. For example, low variability in spot eccentricity (ecc_std) supported assignment to P1 or P2, while trajectories with a high fractal dimension (FractalDim), a measure of how densely a trajectory fills space, were associated with P2. The extent of motion along the short cell axis (track_pc1_sd) was a defining feature of P3, consistent with free diffusion across the full cell width, while P4 could not be assigned from mobility and cellular position alone, but showed a strong signature of high spot eccentricity, above MSD and D_app_.

SHAP further allowed feature interactions to be quantified systematically (Fig. 4E; Supplementary Fig. 4). For example, P1 had the strongest pairwise interaction between D_app_ and distance to the nucleoid. Low D_app_ increased the probability of P1 assignment for trajectories outside the nucleoid, while this effect was largely lost for trajectories inside or near the nucleoid (Fig. 4F). This also explains how the cellular context resolves molecular behaviour beyond motion alone. Because each population contained trajectories from several proteins, we next asked how the molecules of individual proteins partition among these phenotypes.

### Protein-specific occupancy of molecular phenotypes

Rather than assigning each protein to an average class, we quantified the fraction of its trajectories in each molecular phenotype (Fig. 5A). This occupancy profile preserves the heterogeneous behaviour of individual proteins and describes how their molecules partition between modes of motion and cellular environments.

**Fig. 5.**
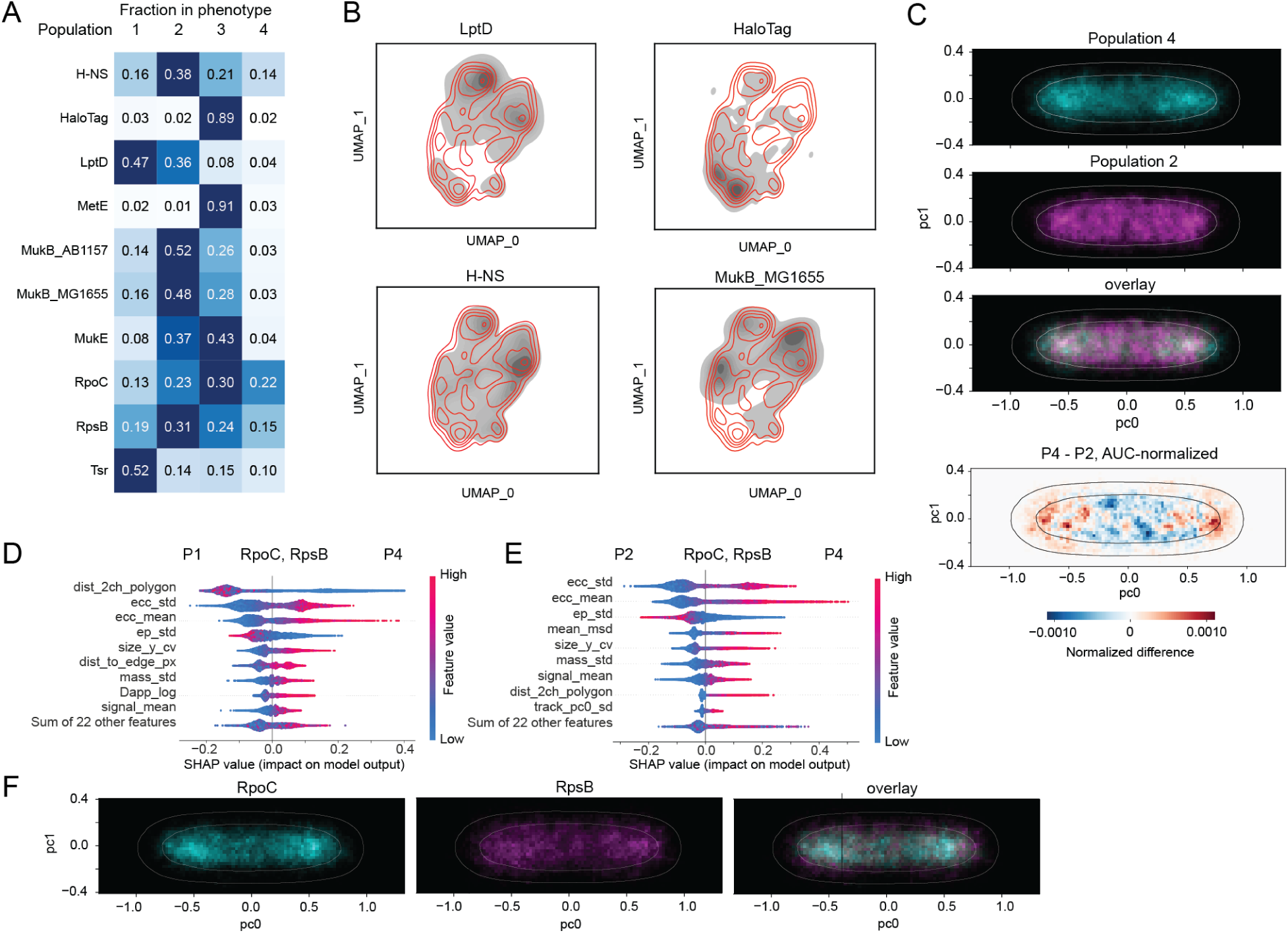
Occupancy profiles quantify protein behaviour and reveal transcription-translation phenotype at the nucleoid periphery. **A**, Fraction of each protein’s trajectories assigned to populations P1–P4; unassigned and artefact trajectories are omitted. Proteins with simple expected behaviour show skewed profiles: Tsr (0.52) and LptD (0.47) occupy mainly the membrane-associated P1, and HaloTag (0.89) and MetE (0.91) the freely diffusing P3. MukB (0.48–0.52) and H-NS (0.38) are enriched in the nucleoid-associated P2 but also occupy the other populations, whereas RpoC and RpsB occupy all four populations, with shared enrichment in P4 (0.22 and 0.15, respectively). **B**, Trajectory density (grey) of LptD, HaloTag, H-NS and MukB_MG1655 in the UMAP embedding over isolines of the reference density (red). **C**, Localization density of RpoC and RpsB trajectories assigned to P4 (cyan) and P2 (magenta) in normalized cell coordinates, their overlay, and the difference between AUC-normalized density maps (P4 − P2; red, enriched in P4; blue, enriched in P2). Lines, dataset-wide cell and nucleoid outlines. P4 localizations are enriched at the nucleoid periphery and P2 localizations within the nucleoid. **D-E**, SHAP summary plots for binary GBC separating P4 from P1 (**D**) and P4 from P2 (**E**) for RpoC and RpsB trajectories. Distance to the nucleoid separates P4 from the membrane-associated population, whereas emitter properties (eccentricity mean and s.d., static error s.d.), separate P4 from the nucleoid-associated population. **F**, Localization density of RpoC (cyan) and RpsB (magenta) trajectories in P4, and their overlay. Both proteins show a similar granular pattern around the nucleoid. The composition, intermediate mobility, peripheral position and emitter properties of P4 are consistent with a molecular environment at the interface between transcription and translation.

Proteins with simple expected behaviours displayed strongly skewed occupancy profiles. The chemoreceptor Tsr, an inner membrane protein enriched at cell poles^45,46^, mapped predominantly to the membrane-associated phenotype P1 (52%, Fig. 2D, 5A). Similarly, LptD, an outer membrane β-barrel protein that translocates lipopolysaccharides to the cell surface^47^, was most frequently assigned to P1 (47%; Fig 5A, B). 36% of LptD trajectories were assigned to the nucleoid-associated phenotype P2, presumably because membrane-bound molecules on the upper and lower cell surfaces project onto the nucleoid in two-dimensional imaging. The cytoplasmic methionine synthase MetE^48,49^, whose interactions are unlikely to substantially alter its mobility or localization, fell almost entirely into the freely diffusing phenotype P3 (91%), as did HaloTag alone (89%; Fig. 5A, B), supporting the validity of the phenotype assignments.

Proteins associated with the chromosome showed broader occupancy profiles. MukB, the SMC homolog of the MukBEF complex, was enriched in P2 (48% in MG1655 and 52% in AB1157), with most remaining trajectories in P3 (28% and 26%, respectively; Fig. 5A, B). MukE was distributed between P2 (37%) and P3 (43%), and the nucleoid-associated protein H-NS^50^ was most frequently assigned to P2 (38%) but was detected in all other phenotypes (Fig. 5A, B). This heterogeneity presumably reflects the partitioning of these proteins between bound and unbound states and between different molecular interactions of the chromosome.

Components of the gene expression machinery showed the broadest distributions. RpsB, a protein of the 30S ribosomal subunit, partitioned between P1 (19%), P2 (31%) P3 (24%), and in P4 (15%), and RpoC, the β′ subunit of RNA polymerase, showed a similarly broad distribution (30%, 23%, 13% and 22%, respectively; Fig. 2D, 5A). Thus, a single protein can occupy several distinct regions of the molecular phenotype landscape.

These analyses show that MIST describes protein behavior as occupancy across robust and interpretable molecular phenotypes. Three of these phenotypes corresponded to well-defined behaviors, while the identity of P4 remained open. Its occupancy profile, with the strongest enrichment of RpoC and RpsB, suggested that it may reflect a specific cellular process.

### MIST identifies a shared phenotype of RNA polymerase and ribosomes *de novo*

Three of these phenotypes corresponded to well-defined behaviours, while P4 emerged from density-based clustering without prior biological specification of the state. P4 was most frequently occupied by RpoC (22% of trajectories) and RpsB (15%) (Fig. 5A), and while both proteins were distributed across several phenotypes, their shared enrichment in P4 distinguished it from the other proteins. The mobility of P4 trajectories was intermediate between P2 and P3 (median D_app_ = 0.33 µm²/s; Fig. 4B), and the D_app_ distributions of RpoC and RpsB overlapped extensively between phenotypes. In contrast, its spatial context was distinct: P4 trajectories were enriched toward the nucleoid boundary, especially towards the cell poles, in comparison to P2 (Fig. 5C).

To identify the features that separate it, we trained binary classifiers on RpoC and RpsB P4 trajectories and compared it with P1 and with P2. Distance to the nucleoid was the dominant feature separating P4 from P1 (Fig. 5D). In contrast, P4 and P2 were distinguished mainly by spot eccentricity and other emitter properties, whereas MSD and distance to nucleoid contributed less (Fig. 5E). Within P4, RpoC and RpsB localizations showed similar distributions with slightly reduced density at mid-cell (Fig. 5F). Together with its localization at the nucleoid periphery (Fig. 5C), this suggests that P4 represents a molecular environment at the spatial interface between transcription and translation. This recapitulates known features of ribosome and RNA polymerase organization. Free 30S subunits are not excluded from the nucleoid, unlike translating ribosomes, and were estimated to comprise 13% of RpsB^51^ (here, 24% of RpsB trajectories in P3). Nucleoid exclusion of mRNA-bound ribosomes or polyribosomes has been attributed to entropic effects of the condensed chromosome^52^, and a similar spatial organization has been reported for RNA polymerase^53^. Using a heterologous T7 promoter, Yang et al.^54^ visualized ribosome assembly within the nucleoid and relocalisation upon translation, and Kim et al.^55^ further showed that this process depends on ribosome-binding-site strength and RNA decay. Transcription-translation translocation does not require direct physical coupling between RNA polymerase and ribosomes.

Notably, while the previous studies were designed to test specific hypotheses and characterized either the gene locus, RNA polymerase or ribosomes, MIST recovered this phenotype for both macromolecules directly from the data *de novo* and defined it not only by co-localization but also by shared motion and emitter properties.

### MIST detects phenotypic changes in novel MukB mutants

A general molecular phenotyping framework should allow new perturbations to be interpreted relative to a fixed reference, without the need to reconstruct the representation. We next tested whether this representation could detect previously unseen functional changes within a single protein.

For this, we analysed five MukB mutants carrying substitutions in ATPase domain (Fig. 6A). In ABC type ATPases, ATP is sandwiched between the Walker motifs of one head and the signature motif of the other, and substitutions in the signature motif or the D-loop impair ATP hydrolysis or ATP-dependent dimerization of the nucleotide-binding domains^56–58^. Two previously characterized mutants, MukB^E1407Q^ and MukB^D1406A^, carry substitutions in the Walker B motif that strongly impair ATP hydrolysis or binding, respectively^59–61^. The three other mutants carry substitutions that have not been characterized in MukB: two in the D-loop, MukB^D1412E^ and MukB^D1412A^, and one in the signature motif, MukB^L1365V,G1368S^. These motifs contribute to ATP-dependent engagement of the two head domains in other SMC proteins^58,62^. All five mutants displayed hallmarks of defective MukB function of temperature sensitivity (Supplementary Fig. 5;^59,60^, establishing that the new D-loop and signature motif mutations also impair MukB function *in vivo*.

**Fig. 6.**
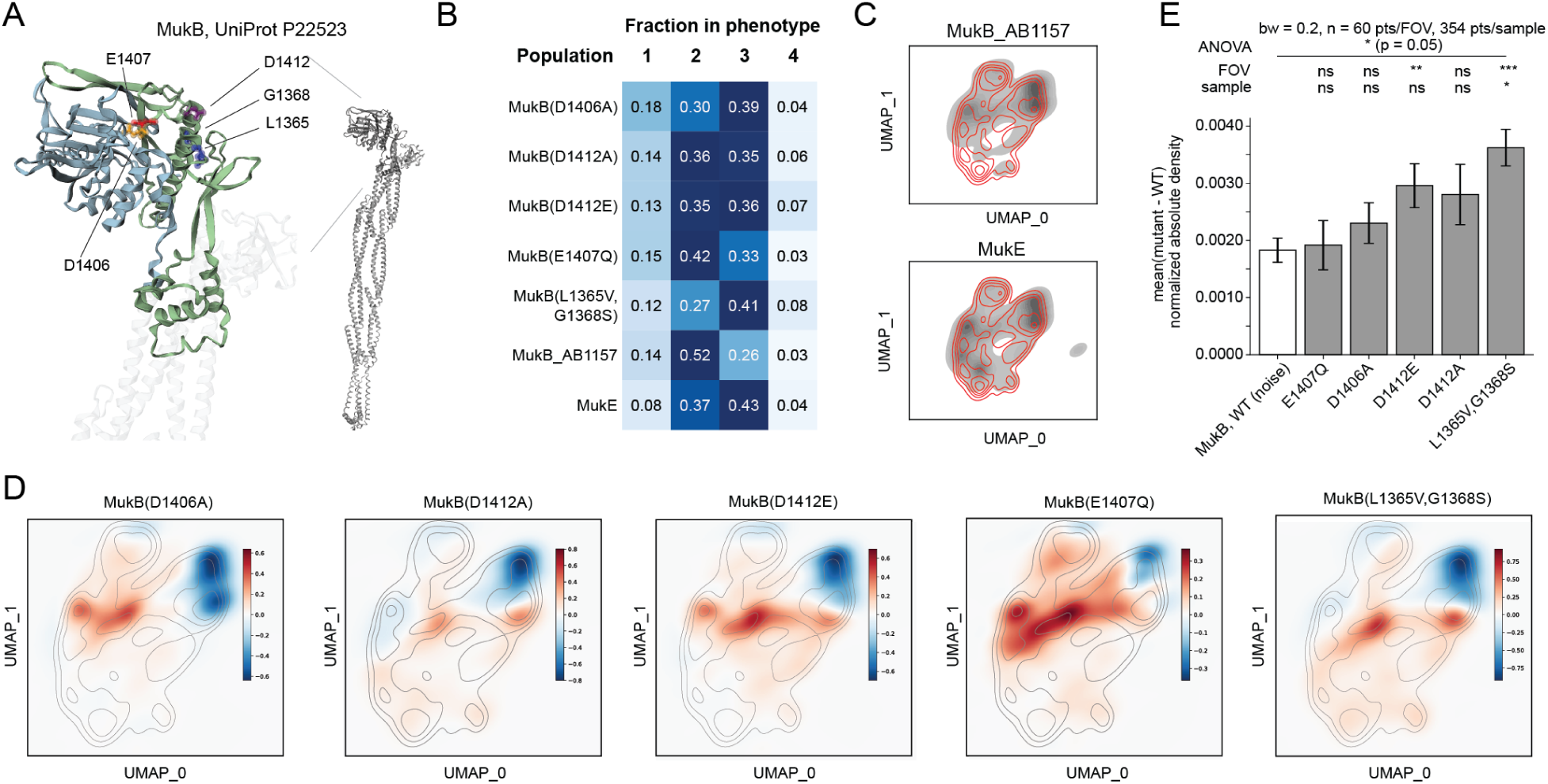
MIST representation detects phenotype redistribution in unseen MukB mutants. **A**, Structural model of E. coli MukB (UniProt P22523), with the ATPase domain enlarged and the substituted residues indicated: D1406 and E1407 (Walker A/B), D1412 (Walker B/D-loop) and L1365 and G1368 (signature motif). **B**, Occupancy of populations P1–P4 for MukB variants, wild-type MukB(AB1157) and MukE. Mutant trajectories were projected into the frozen reference embedding without retraining and assigned to populations by nearest neighbour in the embedding. All variants are depleted in the nucleoid-associated P2 relative to wild type (0.52), most strongly L1365V,G1368S (0.27) and D1406A (0.30), and enriched in the freely diffusing P3. **C**, Trajectory density of MukB(AB1157) and MukE (grey) over reference isolines (red). **D**, Difference in trajectory density between each mutant and wild-type MukB(AB1157) in the UMAP embedding (Gaussian kernel density estimate, bandwidth 0.2; red, enriched in mutant; blue, depleted). Grey lines, reference density isolines. **E**, Perturbation strength, defined as the mean absolute difference between AUC-normalized trajectory density maps of each mutant and pooled wild type (bandwidth 0.2; pixels below 1% of maximum density excluded). Values were calculated per field of view (FOV) and per biological replicate (sample). WT (noise), each wild-type replicate compared with the remaining wild-type replicates. Bars, mean ± s.e.m. of n = 3 biological replicates; n ≥ 9 FOVs. One-way ANOVA across groups at replicate level, P = 0.0499; one-sided Dunnett’s test against WT (noise) at FOV and replicate levels: *P < 0.05, **P < 0.01, ***P < 0.001; ns, not significant. Perturbation strength ranges from close to wild type (E1407Q) to extensive redistribution (L1365V,G1368S or D1412E).

Centroid distances ranked the mutants by the magnitude of their effect but did not reveal which aspects of MukB behaviour changed (Supplementary Fig. 6). Meanwhile occupancy of MIST phenotypes show that all five mutants shifted from P2 toward P3, but to different extents in comparison to wild type (P2 52%, P3 26%). E1407Q caused the smallest redistribution (P2, 42%), whereas the neighbouring D1406A substitution depleted P2 to 30%. The D-loop mutants D1412E and D1412A showed intermediate depletion (35% and 36%), and the signature motif mutant L1365V,G1368S the strongest (27%). The D-loop and signature motif mutants also showed a small increase in P4. In addition, D1406A was close to wild type at the protein level (1.5) yet showed the second-strongest depletion of P2, demonstrating that similar average representations can hide substantial redistribution of individual molecules between phenotypes.

We also compared the full trajectory density landscapes of each mutant and wild-type MukB in the reference embedding (Fig. 6D). Difference maps showed loss of density in the region occupied by nucleoid-associated MukB and gain in the more mobile central region of the landscape. To quantify perturbation strength of each mutant, we calculated the mean absolute difference between the mutant and wild-type density landscapes and normalized it with wild-type replicate variation (Fig. 6E). This analysis revealed a range of perturbation strengths, ranging from E1407Q, close to wild type, to L1365V,G1368S, which produces significant redistribution of MukB molecules (Fig. 6E). E1407Q having the mildest effect may fit a model where ATP binding and head engagement, but not hydrolysis, are needed for chromosome association^56,59,60^. In addition to redistribution among the reference phenotypes, some mutant trajectories occupied a sparsely populated region of the reference representation associated with increased cell area. Together, these results show that the MIST reference space can be used to detect and quantify the molecular consequences of previously unseen genetic perturbations within a single protein.

## Discussion

The dynamics of proteins in living cells are shaped by their interactions and cellular environment, yet most single-molecule analyses quantify them through a single measurement modality and treat cellular context as unwanted variation. Here, we instead treated cellular context as complementary information. MIST integrates trajectory dynamics, emitter properties, cellular context and other measurement channels into a common representation at single-trajectory resolution. Multimodal measurements allow better inference of how the cell state affects protein behaviour. Using a reference panel of nine proteins, we showed that this multimodal description more than doubles protein-specific information relative to trajectory features alone. Data-driven clustering of the resulting space identified recurrent molecular phenotypes, including, de novo, a phenotype shared by RNA polymerase and ribosomes at the nucleoid periphery, and detected graded effects of previously uncharacterized MukB mutations.

Single molecule trajectory analysis has become increasingly informative. Stracy et al.^53^ resolved RNA polymerase populations whose behaviour depended on transcriptional state, and diffusional fingerprinting combined engineered features with hidden Markov model-derived states to separate otherwise similar particle populations^15^. DeepSPT replaced the hidden Markov model with an ensemble of U-Nets that assigns a state to each trajectory step^16^. DeepTRACE extended the diffusion feature space with photometric properties and subcellular localization^17^.

Our benchmarking indicates that the information contained in trajectories, rather than the models used to interpret them, limits trajectory-based analysis. Each feature family increased the intrinsic dimension of the data, showing that emitter properties and cellular context add non-redundant variation. In contrast, engineered trajectory descriptors, DeepSPT and a supervised CNN–Transformer model reached classification performance (macro F1, 0.25–0.32) far below the full MIST representation (0.55). Further gains from trajectory modelling alone are therefore likely to be limited unless additional channels are measured. For example, emitter properties, which have only recently begun to be exploited^17^, are a readily available source of such information in every SMT experiment. Beyond these, MIST can incorporate further orthogonal measurement channels as new feature families, whose contribution can be tested with the same intrinsic-dimension and classification analyses. This shifts the main challenge from how trajectories are modelled to which complementary properties should be measured in the same cells.

MIST was inspired by image-based profiling, which pursues broader aims than state inference, from protein function and interactions to perturbation screens and drug discovery^63^. Cell Painting showed that general organelle morphology carries information comparable to that of omics measurements^26^, and representations learned from images of fluorescently labelled proteins supported a wide range of downstream tasks^29^. In both cases, a single representation is applied across protein and perturbation libraries and supports many analyses rather than a single question. MIST brings this concept to single-molecule tracking and, to our knowledge, is the first framework to combine the reusability of representation and cellular profiling with the molecular resolution of SMT. The two approaches are complementary: image-based profiling reports the state of the cell and infers molecular mechanisms indirectly, whereas MIST reports the behaviour of a specific molecule within that cell.

For this purpose, we kept engineered, physically interpretable features at the core of MIST and added learned features only as an optional module. The CNN–Transformer feature extractor demonstrated this modularity but improved classification only modestly (macro F1, 0.57 vs. 0.55) at the cost of interpretability. Explainability is essential for linking high-dimensional profiles to cell biology^64^.SHAP-based attribution identified the features and feature interactions that define each phenotype; for example, the contribution of mobility to the membrane-associated phenotype depended on distance to the nucleoid. Furthermore, describing each protein by its occupancy across phenotypes provides a molecular analogy of the aggregated profiles used in image-based profiling, but preserves the intra molecular heterogeneity that averaging removes.

The transcription-translation phenotype illustrates how such representations can generate hypotheses. Previous studies showed that gene expression-dependent translation induced gene relocalisation with engineered conditional expression constructs^54,55^. Here, we detected similar phenotype for both RNA polymerase and the ribosome in a minimally perturbed system. Importantly, our experiments were not designed to address this question; the phenotype emerged from data-driven analysis. MIST therefore provides a semantically rich representation from which hypotheses can be generated for a wide variety of downstream applications.

Because new samples are projected into a fixed reference space without retraining, MIST enables perturbation profiling at the molecular level, analogous to Cell Painting, in which the effects of new perturbations on cellular morphology are interpreted against a reference compendium. Projecting five MukB mutants, three of them previously uncharacterized, showed not only that their behaviour changed but also how: occupancy profiles revealed a shift from chromosome-associated toward freely diffusing behaviour, and density landscapes quantified perturbation strength against wild-type replicate variation. The effects agreed with the known requirements of the MukB ATPase cycle, with hydrolysis-deficient E1407Q causing the mildest and nucleotide-binding-deficient D1406A a strong redistribution^59,60^, which supports the interpretation of the new D-loop and signature motif mutants. D1406A also illustrates the value of distribution-level phenotyping, as it remained close to wild-type MukB in protein-level averages yet strongly redistributed between phenotypes. Genetic mutations are not the only perturbations accessible in this way; chemical perturbations and environmental changes, such as altered temperature, can be studied against the same reference.

Several limitations should be considered. First, MISTD v0.1 data set is modest in scale compared with cellular profiling compendia^65^: it contains one tagged protein per sample, 31 selected features and two context channels, from one organism and growth condition. We therefore regard the current phenotypes as a first partition of the molecular phenotype space, which can be refined or split as the reference grows. Second, two-dimensional imaging projects molecules on the upper and lower cell surfaces onto the nucleoid, which presumably explains the assignment of 36% of LptD trajectories to the nucleoid-associated phenotype; three-dimensional tracking would reduce such ambiguity^66,67^. Third, emitter properties such as spot eccentricity and intensity depend on the fluorophore, optics, focus and axial position as well as on molecular properties, so transferring a reference space between instruments will require calibration and batch correction, as developed for image-based profiling^25^. Finally, context features are sensitive to genetic background, as shown by the separation of MukB expressed in AB1157 and MG1655, so comparisons require matched strains.

MIST is modular, and each component can be extended independently. The current version describes cellular context through cell and nucleoid organization, but further feature families, such as markers of membranes, RNA or condensates, stains used in Cell Painting, three-dimensional tracking and organelle markers in eukaryotic cells, can be added. We expect that this type of data sets expand across proteins, organisms and laboratories, and they will support pretrained reference models for single-molecule data. We anticipate applications in fundamental research, such as assigning functions to uncharacterized proteins and characterizing responses to environmental conditions, and in applied research, including drug discovery, antibiotic screening and protein engineering. More broadly, MIST extends the profiling paradigm from cells to individual molecules, providing molecular phenotypes that complement cellular profiles and link the organization of the cell to the behaviour of the molecules within it.

## Materials and Methods

### Bacterial strains, media and growth conditions

Plasmids and strains used in this study are listed in Supplementary Table 2, and Supplementary Table 3, respectively. All strains are derivatives of *E. coli* K-12^68^ MG1655 or K-12 AB1157, as indicated in the supplementary information. All strains were verified by PCR and sequencing. All experiments were performed at room temperature (22°C), the permissive temperature for the temperature-sensitive MukB mutants.

For microscopy, cells were inoculated in a fresh overnight culture and grown in M9GlyCAAT medium containing 0.2 % (v/v) glycerol, 1 g/L casamino acids, 1 mg/L thiamine-HCl, 12 mM MgSO_4_, 0.1 mM CaCl_2_, 6 g/L Na_2_HPO_4_, 3 g/L KH_2_PO_4_, 0.5 g/L NaCl and 1 g/L NH_4_Cl. The following day, cells were diluted ∼1000-fold and allowed to grow to optical density (OD600) 0.1 – 0.3. HaloTag was labelled with Janelia Fluor JFX554, as described in Banaz et al.^69^. Briefly, cells were incubated with 1.5 μM of the JFX554 ligand in the growth medium for 30 min with shaking, washed four times with growth medium, and allowed to recover 60 min with shaking. 30 min before the end of the recovery phase, the nucleic acid stain SYTO 9 (Thermo Fisher Scientific) was added to a final concentration of 0.5 μM. After recovery, the cells were washed one final time with growth medium and resuspended in fresh medium. Cells were immobilized using 1 % (w/v) agarose pads containing low-gelling agarose (Bio-Rad Laboratories) and M9glyCAAT medium. Agarose solution was pipetted onto a fresh microscope slide, and a coverslip was placed over the agarose pad to ensure a flat and uniform surface. Cells were spotted onto the agarose pad, and the sample was imaged immediately.

### Microscopy

Live cell single-molecule tracking (SMT) was performed on a custom-built total internal reflection fluorescence (TIRF) microscope built around the Rapid Automated Modular Microscope (RAMM) System (ASI Imaging) with a motorized stage, a z-motor objective mount, and a motorized filter wheel (US-2000, MIM4, TG8, ASI Imaging). SYTO 9 fluorescence was measured with a 100 mW 488 nm laser with 5 % transmission (iChrome MLE, Toptica). JFX554 fluorescence was measured with a 100 mW 561 nm laser with 70 % transmission (iChrome MLE, Toptica). The lasers were collimated and focused through a 100x oil immersion objective (CFI Plan Apochromat Lambda D, N.A. 1.45, Nikon) onto the sample using an angle for highly inclined thin illumination (Tokunaga et al., 2008). Fluorescence emission was filtered by a dichroic mirror and notch filter (ZT405/488/561rpc and ZET405/488/561m, Chroma). Fluorescence emission was measured using an EMCCD camera (iXon Life 897, 512x512 pixels, Andor). Transmission illumination was provided by an LED source and condenser (ASI Imaging). SMT movies with 20,000 frames were acquired with a frame time of 5.475 ms. All imaging was conducted at RT.

### Single molecule tracking and feature engineering

Localizations were detected with trackpy^70^ (0.6.2). Resulting data frame was used for manual subpixel registration of the SMT channel to the binary cell masks obtained with Omnipose^71^. Next, localizations were joined into tracks for each cell using trackpy, with the localizations outside Omnipose mask excluded. The nucleoid channel image was registered to the SMT channel and the masks, corrected for neighbouring cell signal spillover, normalized and converted to areas after Otsu thresholding.

For each track, we computed 51 features grouped into four families: tracking, emitter properties (Em), cellular coordinates (PC_coord) and morphology (Morph). Tracking features included the apparent diffusion coefficient, track length, the diffusional fingerprinting set but without the hidden Markov model-derived features^15^, turning-angle statistics and MBR. MBR was calculated, as described in ^72^, with each future displacement projected onto the preceding displacement by dot product to obtain a two-dimensional implementation. Emitter properties (brightness, size, eccentricity, signal and localization error) were extracted directly from the trackpy output and summarized per track by their mean, standard deviation and coefficient of variation. Raw xy coordinates were converted to cellular coordinates by PCA [scikit-learn]^73^. PC_coord features describe track position and spread along the cell axes and the distance to the cell edge. Morphology features comprise cell area, aspect ratio and distance to the nucleoid. For highly correlated features (r > 0.9), only one feature was retained per pair. The remaining features were normalized, cleaned from outliers and transformed using scikit-learn’s RobustScaler, IsolationForest (contamination of 0.1) and Yeo–Johnson transformation, respectively. Variance gain was assessed from feature contributions to the principal components whose explained variance exceeded that of permuted data. Separation power was assessed as the ratio of mean absolute linear discriminant analysis coefficients obtained for strain to those obtained for biological replicates. Removing features with low contribution or low ratio yielded the final set of 31 features.

### Dimensionality reduction and clustering

Pairwise Euclidean distances between mean feature values of each protein were computed with scipy.spatial.distance and clustered hierarchically (average linkage) with seaborn clustermap. The two-dimensional UMAP projection was computed with umap-learn (Euclidean metric, min_dist = 0.005, n_neighbors = 6, spread = 1.6), with parameters chosen after manual hyperparameter search. Unseen data were projected into the trained embedding without refitting. Transformer-derived features were embedded with the same parameters. Intrinsic dimensionality was estimated from minimal neighbourhood information (TwoNN) in scikit-dimension^41,42^. and compared with that of the same data after independent permutation of each feature.

Semantic content in the supervised setting was analysed by training a multilayer perceptron (MLP) from scikit-learn to predict protein identity. The MLP had three hidden layers of 100 units and was trained by stochastic gradient descent with early stopping. Inputs were tracking, transformer-derived and DeepSPT^16^ features, as well as cumulative combinations of the handcrafted feature families. MLP performance was assessed by the multiclass macro-averaged F1 score on the test sets of five random 80:20 cross-validation splits, stratified by protein and biological replicate. HDBSCAN was used for clustering, with min_cluster_size=3000, min_samples=15. KNN majority vote by KNeighborsClassifier (scikit-learn, k = 5) was used to expand HDBSCAN-defined clusters. Populations isolated by clustering were used to train gradient boosting classifiers (GBC; scikit-learn), one class against all and for selected pairs of proteins or populations. Logistic regression served as a baseline, and both were evaluated by balanced accuracy on 20% of tracks held out from training. Trained GBCs were then explained by SHAP. SHAP interactions were approximated by shap.utils.approximate_interactions and symmetrized by averaging the interaction matrix with its transpose.

### Deep learning

Each trajectory was converted into 10 input channels: x and y displacement between frames, step length, log-transformed step length (calculated as ln(1 + s_t), where s_t is the step length at time step t), squared displacement between the positions at time steps t and t − 2, smoothed step length (the sum of the current step length and the previous two step lengths, divided by three), movement direction (scalar angle in radians), turning angle calculated as the following step’s direction angle minus the current step’s direction angle and wrapped to [−π, π), and centred x and y coordinates (calculated by subtracting the mean x and y coordinates from the corresponding coordinates). The means were calculated separately for each trajectory using only valid, non-padded time points. These input features were calculated from input two-dimensional coordinates in the model. The values at padded steps were set to zero.

The input was processed by two parallel one-dimensional convolution layers with kernel sizes of 3 and 7. Each convolutional layer produced half of the embedding channels. The output of each layer was passed through layer normalization and GELU activation. The outputs were first concatenated along the channel dimension and then combined by a 1 × 1 convolution layer, also followed by layer normalization and GELU activation, plus dropout. A residual path was formed using a separate 1 × 1 projection and layer normalization. The outputs of the residual and CNN paths were added, and the padded positions were set to zero. Related combinations of convolutional and Transformer layers have been used to characterize diffusion trajectories^74^.

A learnable classification [CLS] token^75^ was prepended to the CNN outputs, and the sinusoidal positional encodings^76^ were added to the results. The Transformer used multi-head self-attention and feedforward layers with GELU activation. Layer normalization was applied to the input of each self-attention and feedforward sublayer, following the Pre-LayerNorm formulation^77^. A separate layer normalization was applied after the encoder stack. A key-padding mask prevented attention to padded positions, while the [CLS] token remained unmasked.

The final [CLS] representation was projected to 16 dimensions by a linear layer and then normalized by layer normalization. This output served as the trajectory features and the input to the classifier. The classifier was a multilayer perceptron with three fully connected hidden layers of 100 units each, and each hidden layer was followed by a ReLU activation. The final linear layer produced ten logits. During training, these logits were passed directly to the cross-entropy loss. During inference, softmax was applied to the logits to obtain class probabilities.

This model was implemented in PyTorch^78^. Hyperparameter search was completed using Optuna ^79^ to maximize the best validation macro F1 score achieved during each trial. The search space included the number of attention heads, the dimension of each head, the encoder depth, the feedforward width, the dropout rate, the learning rate, and the weight decay. The embedding dimension was the product of the number of heads and the dimension of each head.

## Data, Code and Material availability

Strains and plasmids are available upon request. Data will be deposited to a repository. The analysis code is available at: https://github.com/Makelalab/Multimodal-Interpretation-of-Single-molecule-Trajectories-MIST

## Author contributions

Conceptualization, T.R. and J.M.; Methodology, T.R., L.R., Y.S.; Investigation, T.R., L.R., Y.S. and J.M.; Funding acquisition, J.M.; Formal analysis, T.R., L.R., Y.S.; Data curation T.R., L.R., L.R.; Visualization T.R., L.R., Y.S.; Project administration, J.M; Software T.R., Y.S.; Supervision, J.M; Writing – original draft, T.R. and J.M.; and Writing – review & editing, T.R., L.R., Y.S. and J.M.

## Acknowledgements

We thank other members of the Mäkelä group for their support, discussion, and critical reading of the manuscript. We thank Noora Roos for the H-NS-HaloTag strain. We are grateful for gift of JFX554 from Dr. Luke Lavis. We also thank Stephan Uphoff for the gift of pBAD-Halo and sending the strains. This work was supported by the European Research Council (ERC StG TEMPADAPT, 101075984) and the Research Council of Finland (349116 and 353572). The computations and data handling were enabled by resources provided by the CSC – IT Center for Science. The funders had no role in study design, data collection and interpretation, or the decision to submit the work for publication.

## Supplementary figures and legends

**Supplementary Fig. 1.**
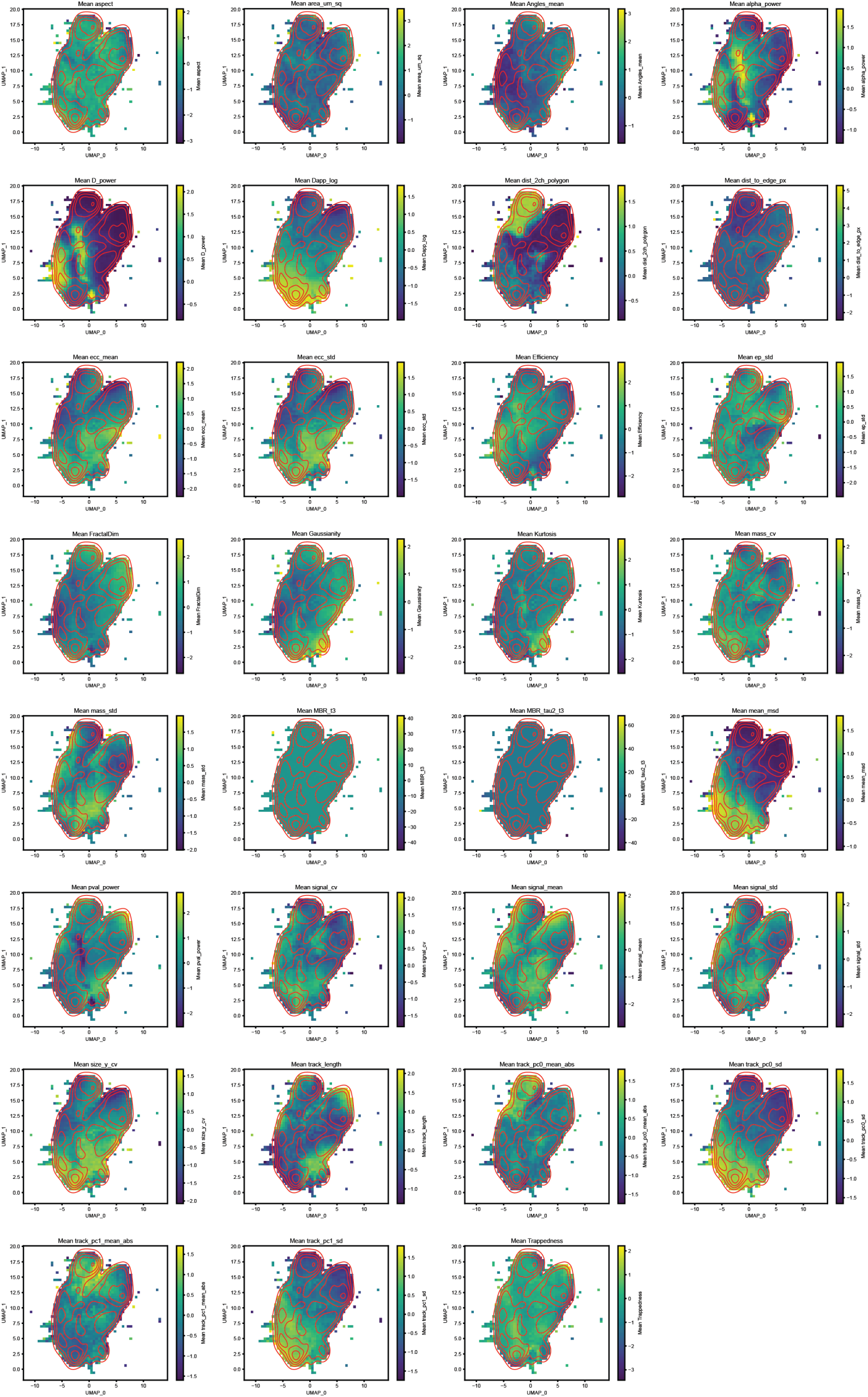
Feature maps of the MIST phenotype space. Mean transformed and standardized value of each of the MIST features per bin of the UMAP embedding of reference trajectories (Fig. 2b). Red lines, five linearly spaced isolines of trajectory density, identical in all panels. Features vary non-continuously across the embedding, and no single feature accounts for its structure, which reflects combinations of motion, emitter and cellular-context features.

**Supplementary Fig. 2.**
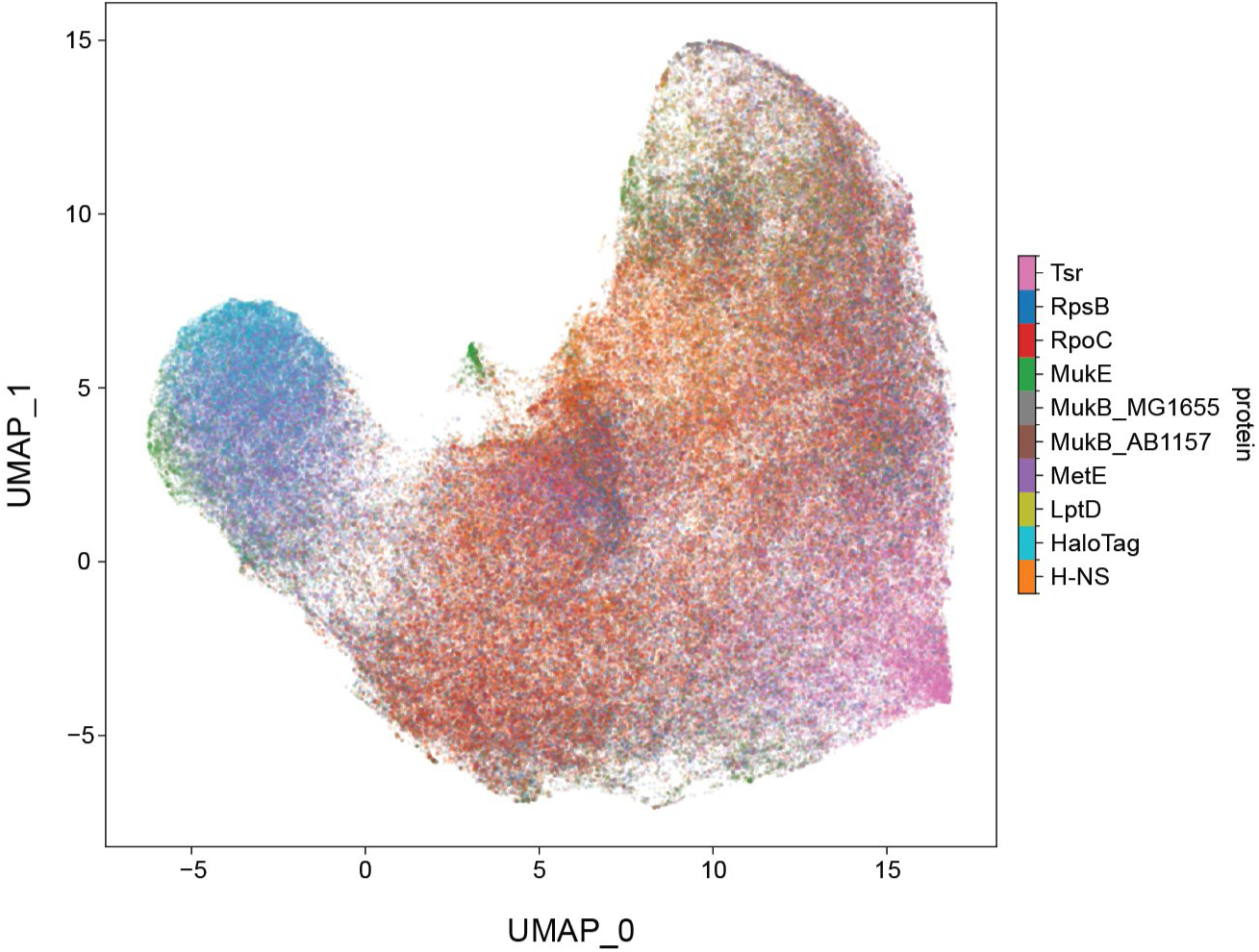
UMAP projection of the learned DL latent space. Each point is one trajectory, coloured by protein. Proteins occupy more separated regions than in the MIST embedding, but the large-scale organization of the embedding does not reflect functional relationships between proteins.

**Supplementary Fig. 3.**
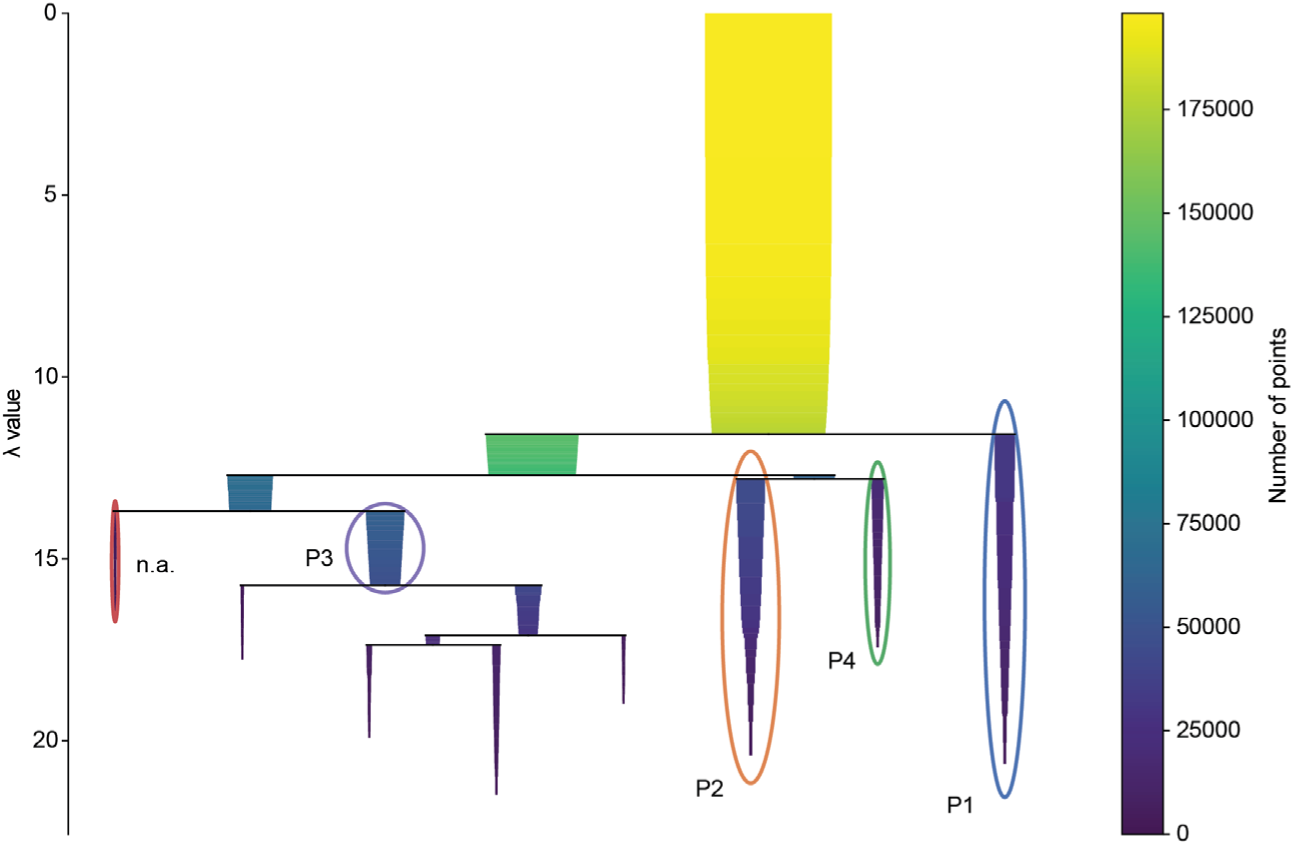
HDBSCAN condensed cluster tree. Condensed tree from HDBSCAN (minimum cluster size 3,000; minimum samples 15) applied to the two-dimensional UMAP embedding of reference trajectories. λ, 1/distance; branch width and colour, number of trajectories. Circled branches, selected clusters. Unassigned trajectories were assigned by 5-nearest-neighbour majority vote unless their mean neighbour distance exceeded the 70th percentile.

**Supplementary Fig. 4.**
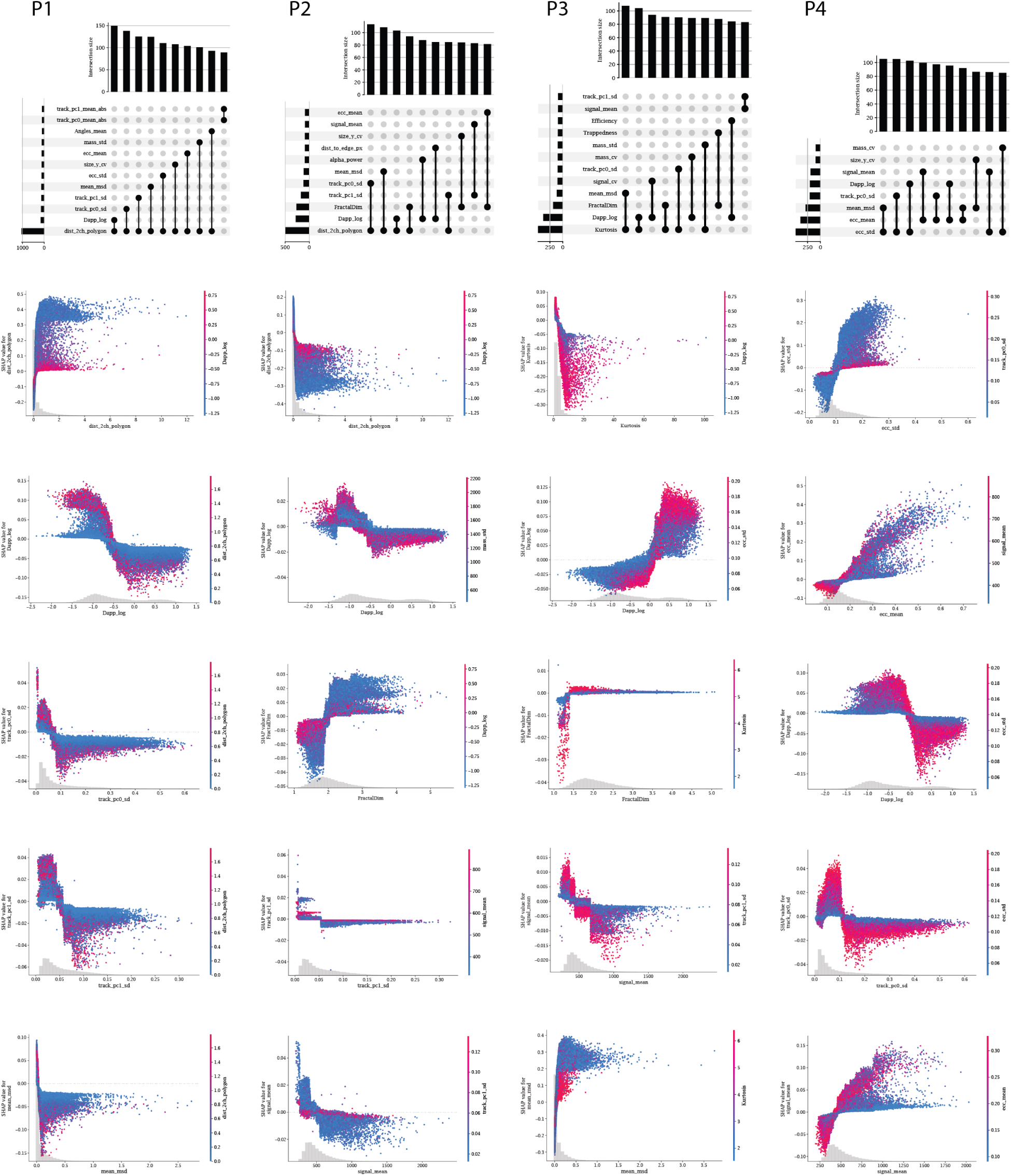
Pairwise SHAP feature interactions for each molecular phenotype. For each population (P1–P4), top: ten strongest pairwise feature interactions from the one-versus-rest gradient-boosting classifiers in Fig. 4d, ranked by approximate SHAP interaction strength; below: SHAP dependence plots for the top-ranked features, each coloured by its strongest interaction partner. Grey histograms, feature value distributions. Distance to the nucleoid dominates the interactions for P1 and P2, whereas interactions for P3 and P4 are dominated by motion and emitter features (kurtosis for P3, eccentricity for P4).

**Supplementary Fig. 5.**
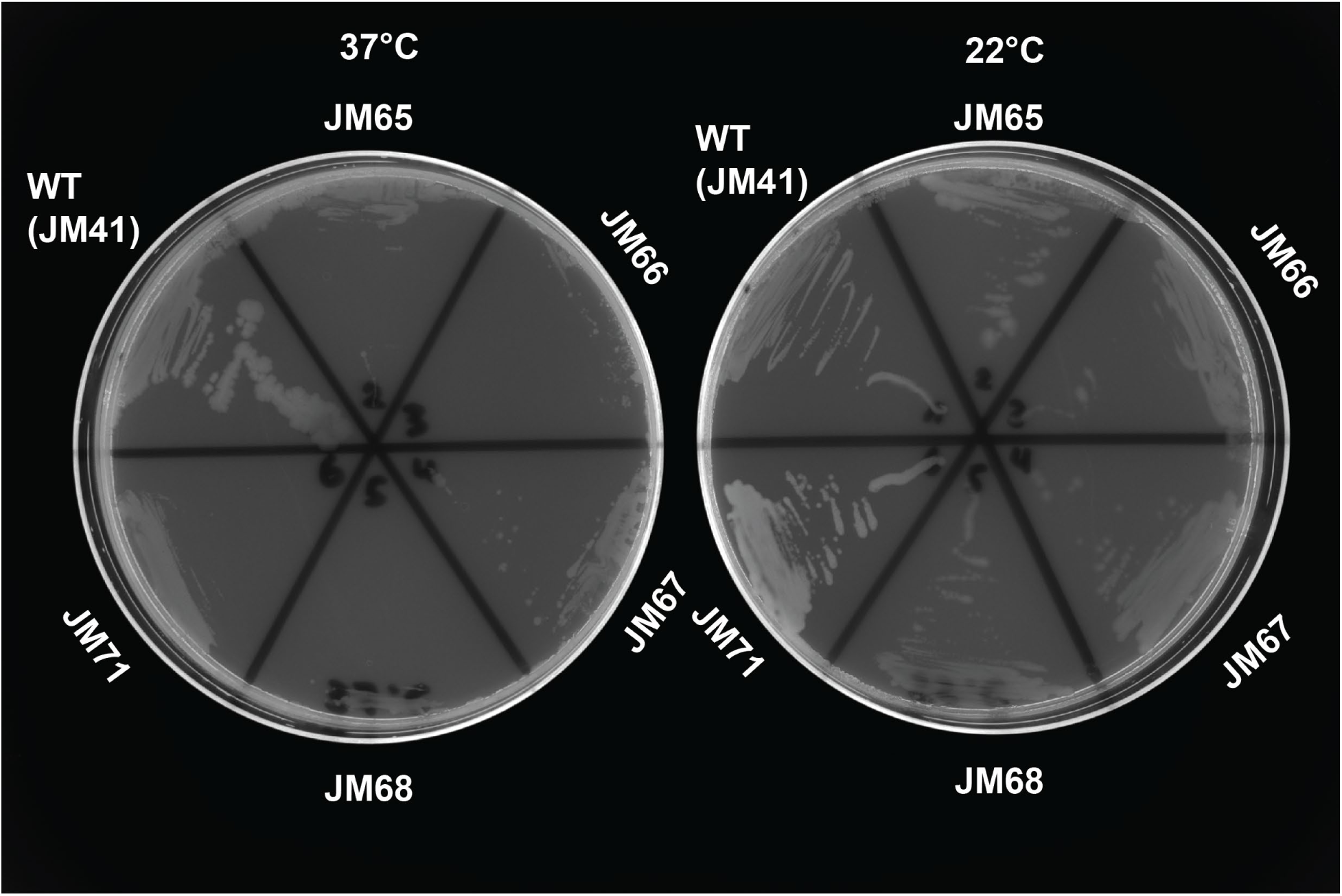
Temperature sensitivity of MukB mutants. Strain plated at 37°C and 22°C. Strain listed in Supplementary Table 3.

**Supplementary Fig. 6.**
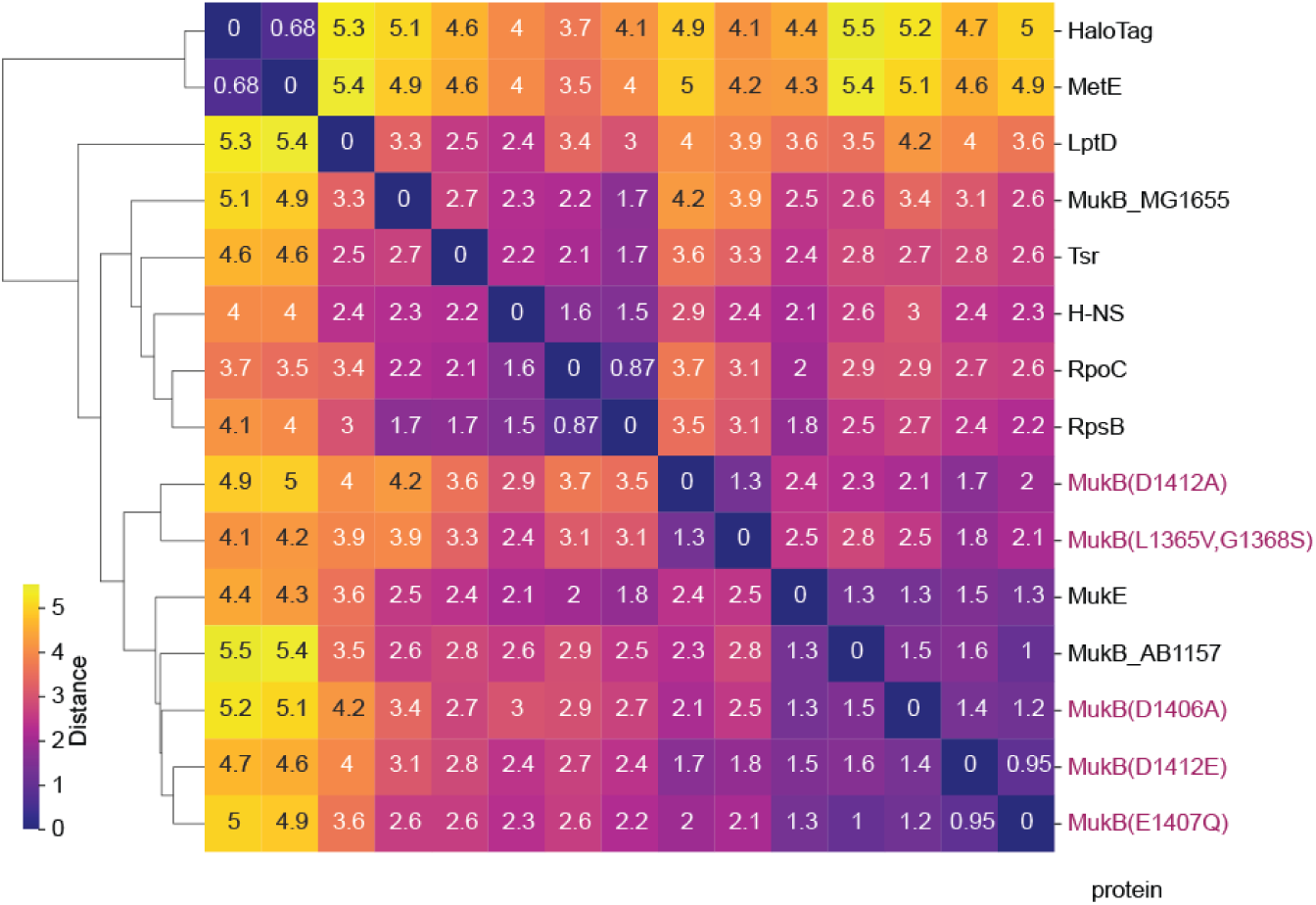
Protein-level distances between reference proteins and MukB mutants. Euclidean distances between centroids of reference proteins and MukB variants (magenta font) in the MIST feature space, with average-linkage hierarchical clustering. Mutant trajectories were transformed with models fit on the reference dataset only. E1407Q, D1406A and D1412E remain close to wild-type MukB(AB1157, 1.0–1.6) and cluster with it and MukE, whereas D1412A and L1365V,G1368S form a neighbouring branch (2.3 and 2.8 from wild type). Protein-level averages capture only part of the redistribution detected by phenotype occupancy (Fig. 6).

**Supplementary Table 1.** Descriptive statistics of particles in different protein states as defined by density-based clustering.

| Cluster N / Phenotype | Variable | Mean | Median | Q25 | Q75 |
|---|---|---|---|---|---|
| 1 / Membrane | Dapp_log | 0.11 | 0.106 | 0.071 | 0.158 |
|  | dist_2ch_polygon | 0.13 | 0.095 | 0.045 | 0.176 |
|  | dist_to_edge_px | 0.17 | 0.166 | 0.109 | 0.233 |
|  | track_pc0_sd | 0.03 | 0.028 | 0.019 | 0.042 |
|  | track_pc1_sd | 0.03 | 0.024 | 0.017 | 0.035 |
| 2 / Nucleoid | Dapp_log | 0.11 | 0.104 | 0.070 | 0.157 |
|  | dist_2ch_polygon | 0.00 | 0.000 | 0.000 | 0.000 |
|  | dist_to_edge_px | 0.29 | 0.298 | 0.234 | 0.353 |
|  | track_pc0_sd | 0.03 | 0.024 | 0.017 | 0.034 |
|  | track_pc1_sd | 0.03 | 0.023 | 0.017 | 0.032 |
| 3 / Free diffusion | Dapp_log | 1.31 | 1.620 | 0.412 | 4.311 |
|  | dist_2ch_polygon | 0.02 | 0.006 | 0.000 | 0.026 |
|  | dist_to_edge_px | 0.28 | 0.284 | 0.244 | 0.321 |
|  | track_pc0_sd | 0.18 | 0.157 | 0.086 | 0.250 |
|  | track_pc1_sd | 0.09 | 0.085 | 0.059 | 0.114 |
| 4 / Transcription-translation coupling | Dapp_log | 0.34 | 0.328 | 0.196 | 0.562 |
|  | dist_2ch_polygon | 0.02 | 0.000 | 0.000 | 0.013 |
|  | dist_to_edge_px | 0.28 | 0.295 | 0.230 | 0.344 |
|  | track_pc0_sd | 0.06 | 0.055 | 0.038 | 0.080 |
|  | track_pc1_sd | 0.05 | 0.044 | 0.030 | 0.062 |

**Supplementary Table 2.**
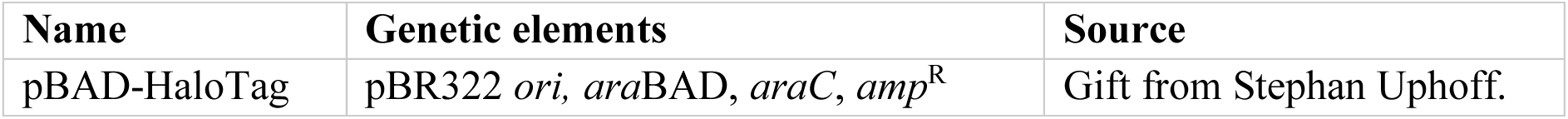
Plasmids used in this study.

| **Name** | **Genetic elements** | **Source** |
|---|---|---|
| pBAD-HaloTag | pBR322 *ori,* *ara*BAD, *araC*, *amp*<sup>R</sup> | Gift from Stephan Uphoff. |

**Supplementary Table 3.** Strains used in this study. *amp* and *kan* refer to plasmid-borne or chromosomal insertions conferring resistance to ampicillin (Amp^r^) or kanamycin (Km^r^), respectively. The chromosomal insertions are flanked by Flp site-specific recombination sites (*frt*) that allow removing of the resistance cassette using a Flp recombinase ^80^. HT refers to HaloTag.

| Name | Background | Genotype | Source |
|---|---|---|---|
| K-12 MG1655 | - | F-, λ-, *ilvG*-, *rfb*-50, *rph*-1 | Mäkelä lab collection |
| CJW7519 | MG1655 | *rpoC*::*rpoC*-HT *frt-kan-frt* | <sup>81</sup> |
| CJW7528 | MG1655 | *rpsB*::*rpsB*-HT *frt-kan-frt* | <sup>81</sup> |
| JM41 | AB1577 | *mukB*::*mukB*-HT *frt-kan-frt* | <sup>69</sup> |
| JM48 | AB1577 | *mukE*::*mukE*-HT *frt-kan-frt* | Mäkelä lab collection |
| JM65 | AB1577 | *mukB*::*mukB*<sup>E1407Q</sup>-HT *frt-kan-frt* | Mäkelä lab collection |
| JM66 | AB1577 | *mukB*::*mukB*<sup>D1406A</sup>-HT *frt-kan-frt* | Mäkelä lab collection |
| JM67 | AB1577 | *mukB*::*mukB*<sup>D1412E</sup>-HT *frt-kan-frt* | Mäkelä lab collection |
| JM68 | AB1577 | *mukB*::*mukB*<sup>D1412A</sup>-HT *frt-kan-frt* | Mäkelä lab collection |
| JM71 | AB1577 | *mukB*::*mukB*<sup>L1365V,G1368S</sup>-HT *frt-kan-frt* | Mäkelä lab collection |
| JML_Ec_01 | MG1655 | *mukB*::*mukB*-HT *frt-kan-frt* | P1 transduction from JM41 to MG1655 |
| JML_Ec_15 | MG1655 | *hns*::*hns*-HT *frt-kan-frt* | Mäkelä lab collection |
| JML_Ec_59 | MG1655 | pBAD-HaloTag | pBAD-HaloTag transformation to MG1655 |
| JML_Ec_136 | MG1655 | *metE*::*metE*-HT *frt-kan-frt* | <sup>11</sup> |
| JML_Ec_173 | MG1655 | *tsr*::*tsr*-HT *frt-kan-frt* | P1 transduction from JM118 (<sup>82</sup>) to MG1655 |
| LT18 | MG1655 | *lptD*::*lptD*-HT-*frt* | <sup>83</sup> |

